# Stimulation of platelet P2Y_1_ receptors by different endogenous nucleotides leads to functional selectivity via biased signalling

**DOI:** 10.1101/2022.08.16.504096

**Authors:** Kate L Arkless, Dingxin Pan, Manu Shankar-Hari, Richard T. Amison, Clive P. Page, Khondaker Miraz Rahman, Simon C. Pitchford

**Author notes:** Author for correspondence and reprint requests: Dr Simon Pitchford, Sackler Institute of Pulmonary Pharmacology, Institute of Pharmaceutical Science, 5.43 Franklin Wilkins Building, 150 Stamford Street, Waterloo Campus, King’s College London, London UK, SE1 9NH. **Authorship:** K.L.A. designed and performed the research, analyzed the data, and wrote the paper. D.P. and R.A. designed research and contributed to project development. K.M.R. designed research and in silico docking analysis and helped write the paper. C.P.P., M.S-H., designed research and helped write the paper. S.C.P. proposed the project, designed research and wrote the paper.

## Abstract

**Background and Purpose:** Platelet function during inflammation is dependent on activation by endogenous nucleotides. Non-canonical signalling via the P2Y_1_ receptor is important for these non-thrombotic functions of platelets. However, apart from ADP, the role of other endogenous nucleotides acting as agonists at P2Y_1_ receptors is unknown. This study compared the effects of ADP, Ap3A, NAD+, ADP-ribose, and Up4A on platelet functions contributing to inflammation or haemostasis.

**Experimental Approach:** Platelets obtained from healthy human volunteers were incubated with ADP, Ap3A, NAD+, ADP-ribose or Up4A, with aggregation and fibrinogen binding measured (examples of function during haemostasis) or before exposure to fMLP to measure platelet chemotaxis (an inflammatory function). *In silico* molecular docking of these nucleotides to the binding pocket of P2Y_1_ receptors was then assessed.

**Key Results:** Platelet aggregation and binding to fibrinogen induced by ADP was not mimicked by NAD+, ADP-ribose, and Up4A. However, these endogenous nucleotides induced P2Y_1_-dependent platelet chemotaxis, an effect that required RhoA and Rac-1 activity, but not canonical PLC activity. Analysis of molecular docking of the P2Y_1_ receptor revealed distinct differences of amino acid interactions and depth of fit within the binding pocket by Ap3A, NAD+, ADP-ribose or Up4A compared to ADP.

**Conclusion and Implications:** Platelet function (aggregation vs motility) can be differentially modulated by biased-agonist activation of P2Y_1_ receptors. This may be due to the character of the ligand-binding pocket interaction. This has implications for future therapeutic strategies aimed to suppress platelet activation during inflammation without affecting haemostasis as is the requirement of current ant-platelet drugs.

## Introduction

The hypothesis of a ‘dichotomy in platelet activation’ was first introduced by Page (1988). In addition to their requisite role in haemostasis, platelets are also important components of the cellular immune system in host defence and in many inflammatory settings. Platelet activation has been shown as an essential precondition for leukocyte activation and recruitment in models of allergic (Pitchford *et al*., 2003; Amison *et al*., 2015; Pan *et al*., 2015) and non-allergic (Kornerup *et al*., 2010; Pan *et al*., 2015; Amison *et al*., 2017) lung inflammation, as well as infection (Youssefian *et al*., 2002; McMorran *et al*., 2009; Amison, O’Shaughnessy, *et al*., 2018). In these settings, platelet activation is not associated with classical aggregation or parameters of primary haemostasis (Amison *et al*., 2015; Cleary *et al*., 2019; Shah *et al*., 2021). Furthermore, we and others have identified platelets in the lungs of patients with asthma, and in animal models of allergic inflammation, sterile inflammation, and infection, suggesting that this cell type can also undergo extravascular migration into sites of inflammation, the antithesis to an aggregatory event (Pitchford *et al*., 2008; Shah et al., 2021; Ortiz-Munoz et al., 2014; Amison, OShaughnessy et al., 2018; Cleary et al., 2020; Le *et al*., 2015). Aberrant platelet activation in the absence of haemostatic dysfunction and thrombosis has also been reported in inflammatory conditions such as rheumatoid arthritis (Boilard *et al*., 2010), acute lung injury (ALI) (Zarbock, Singbartl and Ley, 2006; Looney *et al*., 2009; Grommes *et al*., 2012) and sepsis (Clark *et al*., 2007; Kahn, Hurley and Shannon, 2013; Claushuis *et al*., 2016).

Interestingly, platelet activation in the context of inflammatory function appears distinct from activation involved in haemostasis (aggregation). We have previously provided evidence for biased signalling events at the platelet P2Y_1_ receptor, where we were able to pharmacologically inhibit platelet-mediated leukocyte recruitment, without affecting haemostatic function (Amison *et al*., 2015, 2017). These P2Y_1_-mediated inflammatory responses appear independent of the canonical phospholipase-C (PLC) signalling pathway, instead occurring via Rho-GTPase activity (Amison *et al*., 2015; 2018; Pan et al., 2015), as evidence of an example of biased agonism or ‘functional selectivity’ (Kenakin 2012).

Although P2Y_1_ receptors have been investigated using it’s cognate agonist, adenosine diphosphate (ADP), other endogenous nucleotides are also known to activate this receptor. P3-(5’-adenosyl) triphosphate (Ap3A) has been shown to increase intracellular calcium in 1321N1 cells, downstream of P2Y_1_ receptor activation (Patel *et al*., 2001). Like ADP, Ap3A is stored in platelet dense granules and released upon cell activation (Lüthje and Ogilvie, 1983). However, Ap3A itself is unable to directly elicit platelet aggregation *in vitro*, but can do so indirectly through hydrolysis to ADP via plasma hydrolase activity (Lüthje and Ogilvie, 1984; Lüthje, Baringer and Ogilvie, 1985). Furthermore, Nicotinamide adenine dinucleotide (NAD^+^) causes P2Y_1_ receptor-mediated neuronal hyperpolarisation in *ex vivo* murine studies (Mutafova-Yambolieva *et al*., 2007; Hwang *et al*., 2011). Although no studies that we are aware of have measured intracellular NAD^+^ concentrations within platelets, they do express CD38 on their surface, which is able to hydrolyse NAD^+^ to ADP-ribose (Ramaschi *et al*., 1996; Mutafova-Yambolieva *et al*., 2007). ADP-ribose is another P2Y_1_ receptor agonist, reported to activate this receptor in rat and human primary β-cells through PLC-mediated increases in intracellular calcium concentrations (Gustafsson *et al*., 2011). There is also some evidence to suggest that ADP-ribose may inhibit platelet aggregation (Del Principe *et al*., 1986), potentially by acting as a competitive inhibitor for ADP. Additionally, uridine adenosine tetraphosphate (Up4A) is another endogenous P2Y_1_ receptor agonist, found to elicit relaxation in human and murine colon muscle (Durnin *et al*., 2014), whilst also enhancing vascular contraction in mouse (Zhou *et al*., 2015) and diabetic rat arteries (Mahdi *et al*., 2018). Again, the presence of this agonist has not yet been investigated within platelets, but it is released from endothelial cells upon their activation (Jankowski *et al*., 2005).

Although all of the P2Y_1_ receptor agonists described above have the potential to activate platelets, their effects are incompletely understood. Therefore, the aim of the present study was to evaluate a panel of endogenous P2Y_1_ receptor agonists with respect to platelet activation using haemostatic and inflammatory platelet function assays.

## Materials and Methods

### Materials

Acid Citrate Dextrose (ACD)-A Vacuette tubes (455055) were purchased from Greiner Bio-One. The purinergic receptor agonists, ADP (01905), NAD^+^ (N0632), ADP-ribose (A0752) and Ap3A (D1387), the chemotactic peptide N-formylmethionyl-leucyl-phenylalanine (f-MLP) (F3506) and prostaglandin E_1_ (PGE_1_) were all purchased from Sigma Aldrich. The purinergic receptor agonist, Up4A (BLG-U008-01) was purchased from Enzo Life Sciences. The HTS Transwell 96-well plates (3μm pore size) (10077792) and RPMI 1640 cell media with L-glutamine (12004997) were purchased from Fisher Scientific. The P2Y_1_ antagonist, MRS2500 (2159/1), the P2Y_12_ antagonist, AR-C66096 (3321/1), the PLC inhibitor (U73122), the Rac1 inhibitor (NSC23766) and the Rho-associated kinases (ROCK) inhibitor (GSK429286) were purchased from Bio-Techne.

### Human platelet isolation

For all studies, blood was collected in accordance with local ethical approval from King’s College London (Research Ethics Committee Reference: 10/H0807/99) and adhered to regulations outlined by the Human Tissue Act 2004 as previously described (Amison *et al*., 2018). Blood was collected using ACD-A Vacuette tubes from healthy male and female volunteers who had not taken NSAIDS or other anti-inflammatory drugs in the previous seven days, and who were not prescribed anti-platelet drugs. Whole blood was centrifuged at 133 g for 20 minutes at room temperature. An aliquot of the upper platelet-rich plasma (PRP) layer was then used for aggregation studies *(see below)*. To the remaining PRP, 2.5μM PGE_1_ was added before centrifuging at 800 g for 10 minutes at room temperature. Platelet-poor plasma (PPP) was removed and used for aggregation studies *(see below)*. The platelet pellet was resuspended in RPMI 1640 cell media, again adding 2.5μM PGE_1_ and centrifuging at 800 g for 10 minutes at room temperature. Platelets were then adjusted to a final concentration of 5×10^7^ platelets/ml in RPMI 1640 using an Improved Neubauer chamber (Hawksley & Sons Ltd) for chemotaxis studies *(see below)*.

For fibrinogen binding studies, platelets were isolated via gel-filtration as previously described (Petito *et al*., 2018). Briefly, PRP was added to a Sepharose C12-B column and eluted through using HEPES buffer. Only the cloudy platelet containing media eluted from the base of the column was collected. Platelets were then adjusted to a final concentration of 5×10^7^ platelets/ml.

### In vitro platelet aggregation

The various endogenous purinergic agonists (ADP, Ap3A, NAD^+^, ADP-ribose, and Up4A) were investigated for their effects on platelet aggregation quantified by light transmission aggregometry of stimulated PRP at 595nm at 37ᵒC using a SpectraMax 340PC shaking plate reader (Molecular Devices) as previously described (Amison *et al*., 2018). Briefly, PRP was stimulated with vehicle (PBS) or individual agonists and immediately loaded onto the plate reader. Vehicle stimulated PPP was also used as a control. Measurements were taken at 15-second intervals for 16 minutes under shaking conditions. In some studies, PRP was pre-incubated with vehicle (PBS) or increasing concentrations of the P2Y_1_-specific antagonist, MRS2550, or the P2Y_12_-specific antagonist, AR-C66096, for 10 minutes at room temperature before stimulation with agonists.

### In vitro platelet chemotaxis

Inflammatory platelet function downstream of purinergic receptor activation induced by endogenous agonists was also investigated through *in vitro* platelet chemotaxis, as previously described, but with minor amendments (Amison *et al*., 2018). Washed platelets (5×10^7^/ml) were treated with 2mM CaCl_2_ before stimulation with vehicle (PBS) or individual agonists for 5 minutes at room temperature. In some studies, platelets were incubated with antagonists for 10 minutes at room temperature, prior to agonist stimulation. Platelets (80µl) were then added to the top insert of the 96-well transwell plate, with chemoattractant in the bottom well (0/30nM fMLP in RPMI 1640 cell media). Following 90 minutes incubation at 37ᵒC, media from the bottom chamber was stained with Stromatol (1:0.5) and platelets were quantified using an Improved Neubauer haemocytometer and a Leica DM 2000 LED microscope with an x40 objective lens.

### Molecular docking of ligands with the P2Y_1_ receptor

Molecular docking was performed to generate several distinct binding orientations and binding affinity for each binding mode as previously described (Amison *et al*., 2018). Subsequently, the lowest binding free energy was considered as the most favourable binding mode for the system. AutoDock Smina (Koes *et al*., 2013; Trott and Olson, 2010), which uses the AutoDock Vina scoring function by default, was used for the blind molecular docking of the ligands to the P2Y_1_ structure (protein databank PDB ID: 4XNW,4XNV) for finding the best binding site by exploring all probable binding cavities of the proteins. Smina was performed with default settings, which samples nine ligand conformations using the Vina docking routine of stochastic sampling. Then, Genetic Optimization for Ligand Docking (GOLD) molecular docking was applied for the docking of ADP, Ap3A, NAD, ADP-ribose, and Up4A to the Smina-located best binding site of the P2Y_1_ receptor for performing flexible molecular docking as described elsewhere (Jones *et al*., 1995, 1997). Based on the fitness function scores and ligand binding positions, the best-docked poses for the ligands were selected. The GOLD molecular docking procedure was performed by applying the GOLD suite in the CSD Discovery software (Jones *et al*., 1997). The Genetic algorithm (GA) was used in GOLD ligand docking software to examine thoroughly the ligand conformational flexibility along with the partial flexibility of the protein (Nissink *et al*., 2002). Finally, the 2D ligand-protein interaction map was generated using BIOVIA discovery studio visualiser 2021.

### Statistical analysis and experimental design

Data are expressed as mean ± SEM. Quantification of platelets via microscopy was conducted with the experimenter blinded to the sample identity. Chemotaxis data is normalised to a negative control to give a chemotactic index (CI), due to baseline variations between donors. Group sizes are indicated in figure legends and power calculations were undertaken, dependent on variability of assays based on previous data (Amison *et al*., 2018) or pilot data. In particular, where α-error (degree of significance) is 0.05, and β-error (probability of false error) is 0.1 (90% power), we calculated a sample requirement for aggregation experiments of n= 5-6, and chemotaxis studies of n= 6-8. Data were analysed using GraphPad Prism, with specific statistical tests indicated in figure legends. A *P* value of less than 0.05 was considered significant. An exception was platelet fibrinogen binding studies (Figure 1B, D, F, H, J), where no statistical analysis was made (n=4), because these data have been provided as initial confirmatory evidence to that of platelet aggregation data which was also negative in lack of platelet action (Figure 1A, C, E, G, I). The manuscript complies with the *British Journal of Pharmacology* requirements on experimental design and analysis (Curtis et al., 2018).

**Figure 1.**
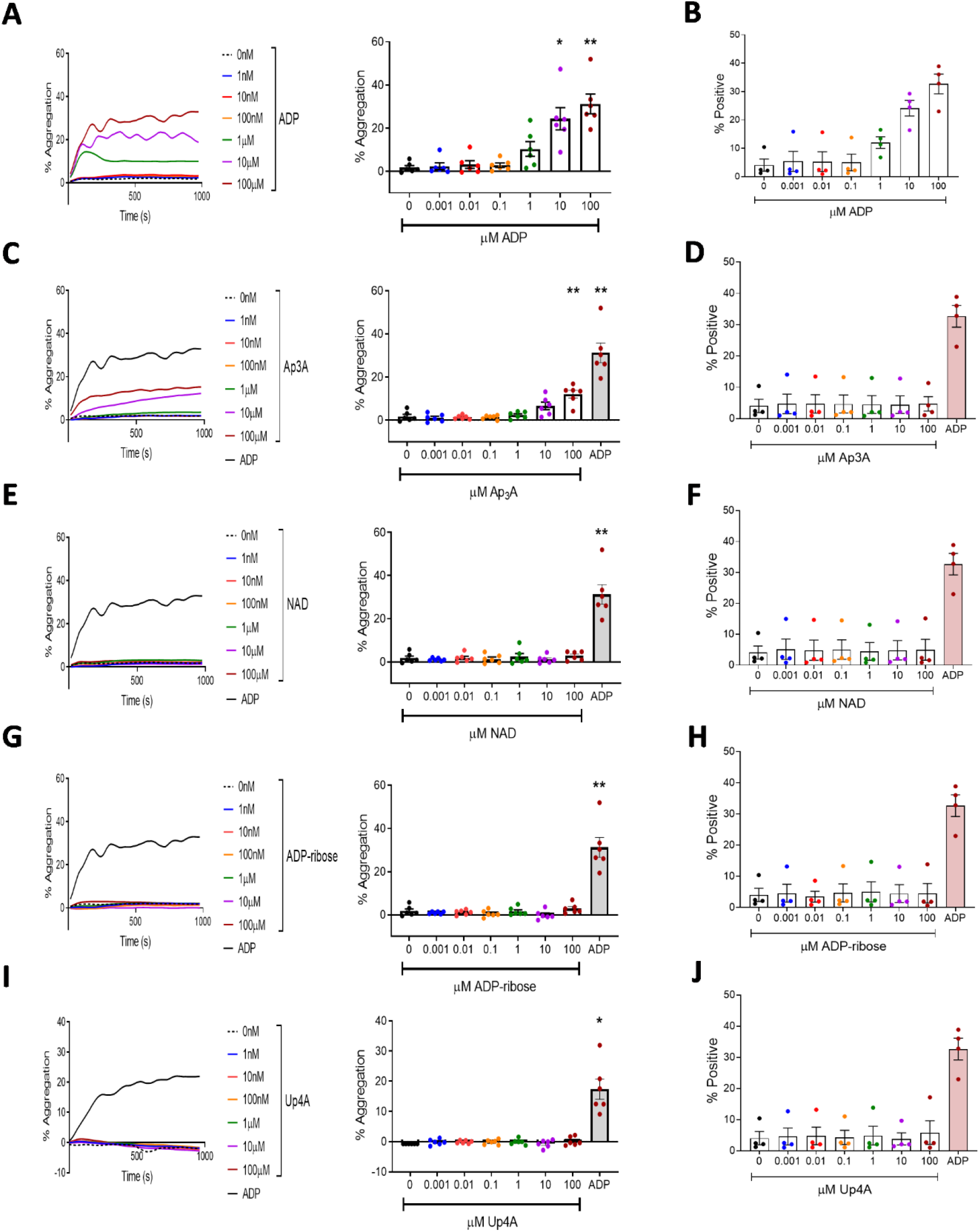
ADP and Ap3A, but not NAD, ADP-ribose or Up4A induce platelet aggregation and fibrinogen binding. PRP isolated from healthy human donors was stimulated with increasing concentrations of endogenous P2Y_1_ receptor ligands, and aggregation measured by light transmission aggregometry, with traces showing affect over 16minutes, and statistical analysis at 5 minutes (A,C,E,G,I). In other experiments, fibrinogen-488 (2µg/ml) was added to gel filtered platelets (0.5×10^8^/ml) and platelets were stimulated with endogenous P2Y_1_ receptor ligands for 30 minutes before fixation and analysis on a Beckman Coulter FC500 to quantify fibrinogen binding (B,D,F,H,J). (A-B) ADP. (C-D) Ap3A. (E-F) NAD. (G-H) ADP-ribose. (I-J) Up4A. Data: Mean ± SEM. n = 6 per group (aggregation), and n = 4 per group (fibrinogen binding). One-way ANOVA with Dunnett’s multiple B) comparisons undertaken on aggregation data only. ^*^P<0.05, ^**^P<0.01 versus negative control (column A).

## Results

### ADP is the only endogenous P2Y1 receptor agonist that induces platelet aggregation

We have confirmed the well established observation that ADP stimulates platelet aggregation via P2Y_1_ and P2Y_12_ receptor stimulation (**Figure S1**). However, in this study, we have also assessed the ability of other endogenous P2Y receptor agonists to elicit this haemostatic platelet function. PRP was stimulated with increasing concentrations of ADP, Ap3A, NAD^+^, ADP-ribose or Up4A. As expected, ADP stimulation led to significant platelet aggregation at both 10µM (P<0.05) and 100µM (P<0.01), with 100µM producing the greatest effect (**Figure 1A**). 100µM Ap3A was also found to elicit significant aggregation (P<0.01) (**Figure 1C**). However, this is likely due to ADP liberation from Ap3A rather than Ap3A itself, as previously described (Lüthje and Ogilvie, 1984; Lüthje, Baringer and Ogilvie, 1985; **Figure S2**). In contrast, neither NAD^+^, ADP-ribose nor Up4A were able to elicit *in vitro* platelet aggregation at any concentration tested (**Figure 1E,G,I**). As an initial confirmatory measure of *in vitro* platelet haemostatic function, a flow cytometric assay to measure platelet fibrinogen binding was optimised to understand possible activation of platelets at the single event level, rather than reliance on the optical density measurement provided by measuring platelet aggregation. 2µg/ml fluorescently labelled fibrinogen (fibrinogen-alexafluor488) was added to gel filtered platelets (0.5×10^8^ platelets/ml) and allowed to acclimatise. Platelets were then stimulated with increasing concentrations of ADP for 30-minutes, fixed using 1% PFA and then analysed on a Beckman Coulter FC500 flow cytometer. With similarity to the platelet aggregation data, only ADP stimulation of platelets resulted in increased fibrinogen binding (**Figure 1A**). Whilst statistical analysis could not be conducted on this data (n=4), none of the other endogenous P2YR agonists caused platelet-fibrinogen binding to occur (**Figure 1B,D,F,H,J**), and was therefore confirmatory to aggregation data.

Given that platelet fibrinogen binding occurs via an integrin αIIbβ3 conformational change, dependent on P2Y_12_ inhibition of adenylyl cyclase cAMP production, it is also suggestive that the endogenous agonists were also not able to cause release of ADP from granular stores, or affect P2Y_12_ activation directly.

### ADP and other endogenous P2Y_1_ agonists contribute to platelet chemotaxis as a measure of inflammatory function

In addition to haemostatic function, the panel of endogenous P2Y receptor agonists was also investigated in the context of inflammatory platelet function. The ability of platelets to migrate to sites of inflammation *in vivo* and to undergo chemotaxis *in vitro* has been shown by multiple groups (Czapiga *et al*., 2005; Kraemer *et al*., 2010; Petito *et al*., 2018) and has been suggested to require P2Y_1_ stimulation (Amison *et al*., 2015; 2018).

Platelets were stimulated with increasing concentrations of ADP, Ap3A, NAD^+^, ADP-ribose or Up4A and chemotaxis towards fMLP was measured using a transwell assay setup. As previously described, platelet stimulation by 100nM ADP led to significant platelet chemotaxis towards fMLP *in vitro* (P<0.0001) (**Figure 2A**) (Amison *et al*., 2018). Like ADP, the P2Y receptor agonist, Ap3A, is also present in platelet dense granules and released into the extracellular milieu upon platelet activation and degranulation (Lüthje and Ogilvie, 1983). However, our results indicate that this endogenous P2Y receptor agonist lacked the ability to elicit platelet chemotaxis towards fMLP (**Figure 2B**). In contrast, NAD^+^, caused *in vitro* platelet chemotaxis towards fMLP at both 1nM (P<0.01) and 10nM (P<0.001), (**Figure 2C**). Similarly, other endogenous P2Y receptor agonists of interest, ADP-ribose and Up4A, also elicited significant platelet chemotaxis *in vitro* expressed as bell-shaped concentration responses. ADP-ribose caused chemotaxis at 1nM (P<0.05), 10nM (P<0.01) and 100nM (P<0.05), with a peak response at 10nM (**Figure 2D**). Finally, Up4A also triggered significant *in vitro* platelet chemotaxis at concentrations of 1nM (P<0.05) and 10nM (P<0.001), with the greatest effect observed at 10nM (**Figure 2E**).

**Figure 2.**
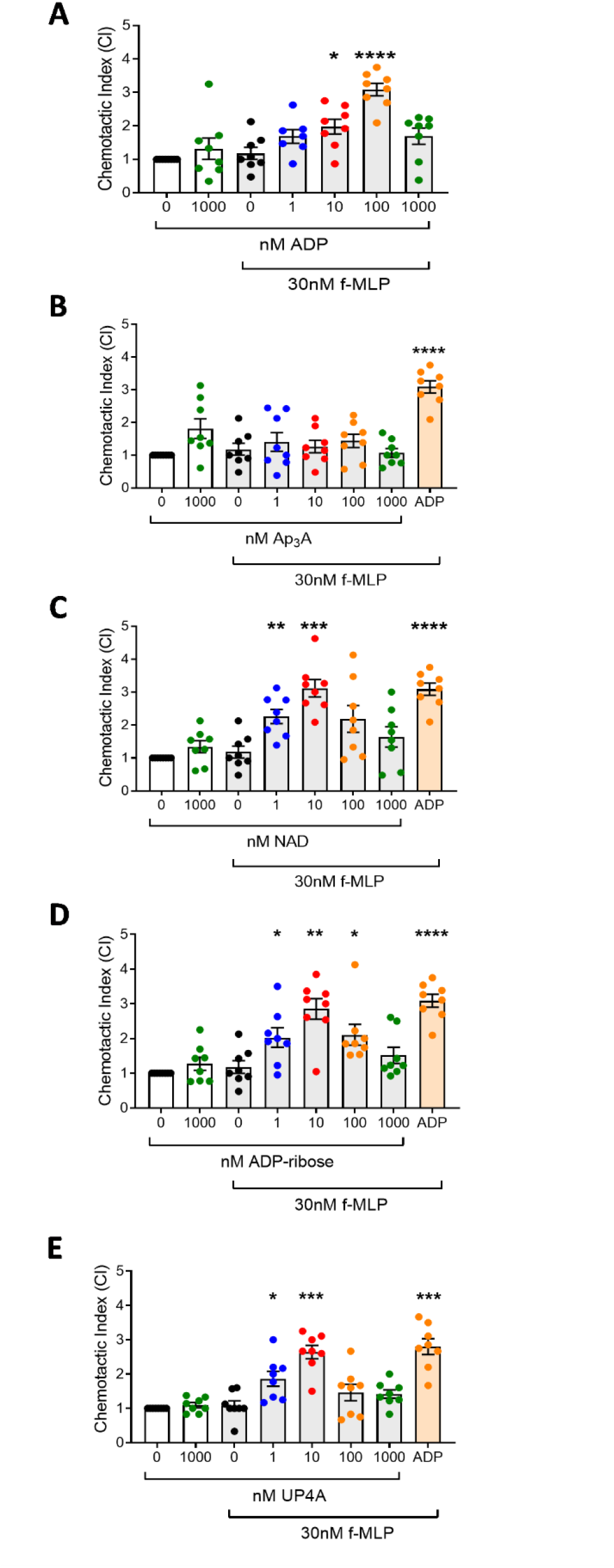
Characterisation of endogenous P2Y_1_ receptor ligands on *in vitro* platelet chemotaxis. Washed platelets (5×10^7^/ml) in RPMI were stimulated with increasing concentrations of endogenous ligands for the P2Y_1_ receptor. Platelets were then added to the top chamber of a 96-well transwell plate, with 0/30nM fMLP in the bottom chamber. After 90 minutes at 37°C, platelets in the bottom chamber were quantified and normalised to negative controls to give the chemotactic index (CI). (**A**) ADP. (**B**) Ap3a. (**C**) NAD. (**D**) ADP-ribose. (**E**) Up4A. Data: Mean ± SEM. n = 8 per group. One-way ANOVA with Dunnett’s multiple comparisons. ^*^P<0.05, ^**^P<0.01, ^***^P<0.001, ^****^P<0.0001 versus in the presence of no nucleotide (column C).

### Endogenous P2Y nucleotides induced platelet chemotaxis via P2Y1 activation, dependent on non-canonical RhoA and Rac1 signalling

In order to understand the role of P2Y receptor activation for endogenous nucleotide induced chemotaxis, we tested this inflammatory function of platelets in the presence of specific P2Y_1_ and P2Y_12_ receptor antagonists. In support of previous studies, we have demonstrated that the P2Y_1_ receptor-specific antagonist, MRS2500, was able to inhibit *in vitro* platelet aggregation induced by ADP (0.1 µM P<0.001, 1µM P<0.001, and 10µM P<0.00001; **Figure 3A**); NAD^+^ (1µM P<0.01, and 10µM P<0.01; **Figure 3C**); ADP-ribose (0.1 µM P<0.01, 1µM P<0.01, and 10µM P<0.01; **Figure 3E**); and Up4A (0.1 µM P<0.05, 1µM P<0.001, and 10µM P<0.01; **Figure 3G**). However, incubation of platelets with the specific P2Y_12_ antagonist AR-C66096 had no effect on platelet chemotaxis requiring stimulation by ADP (**Figure 3B**), ADP-ribose (**Figure 3F**) or Up4A (**Figure 3H**), and whilst AR-C66096 did significantly inhibit NAD^+^-dependent chemotaxis (1µM P<0.01, and 10µM P<0.01), a level of activation remained (**Figure 3D**). These data suggest that the endogenous nucleotides, are similar to ADP, in that they stimulate platelet chemotaxis via P2Y_1_ rather than P2Y_12_ receptor activation.

**Figure 3.**
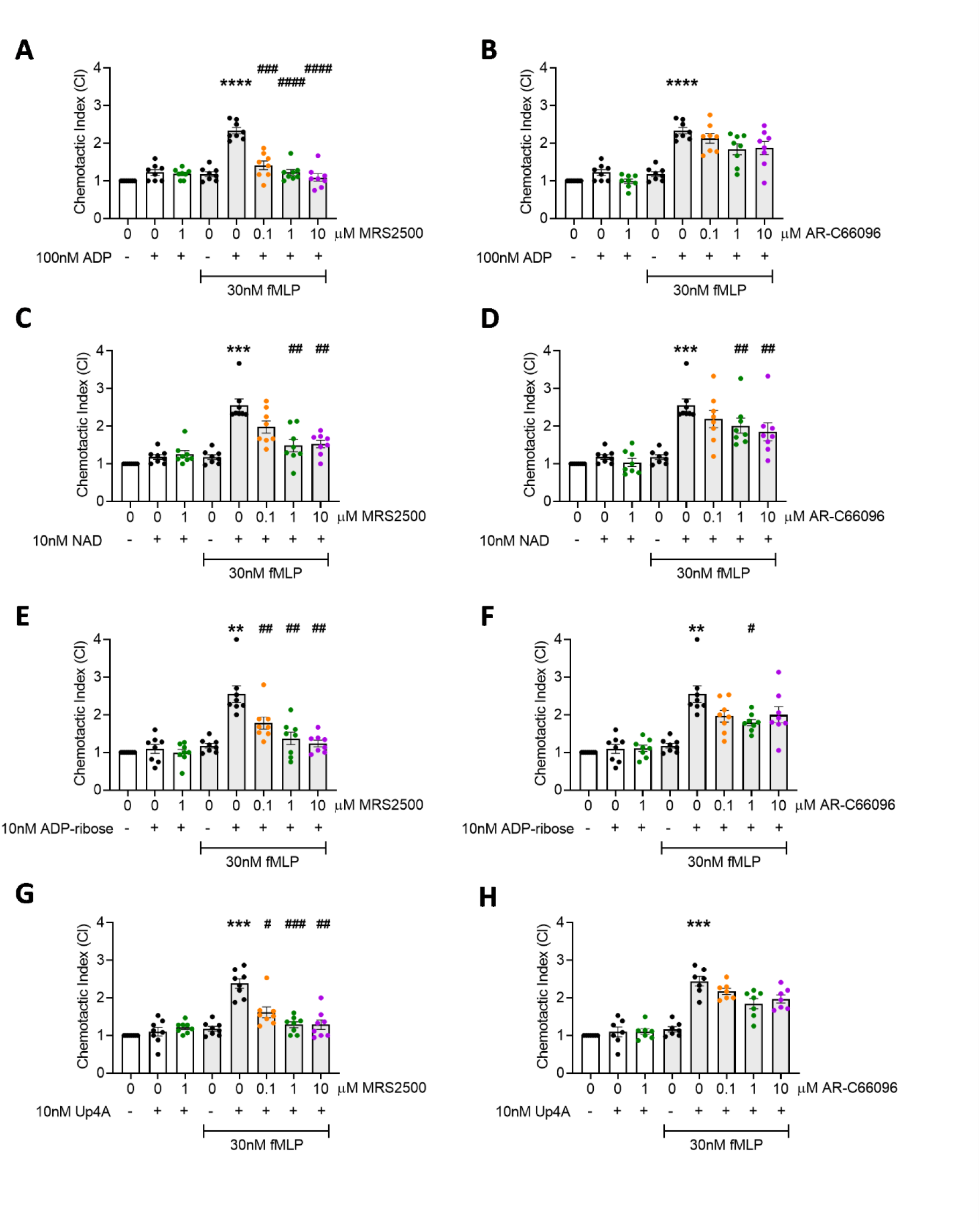
Platelet chemotaxis towards fMLP was induced by ADP, NAD, ADP-ribose, and Up4A via P2Y_1_ receptor. Washed platelets (5×10^7^/ml) in RPMI were incubated with specific antagonists for either P2Y_1_ (MRS2500) or P2Y_12_ (AR-C66096) for 10 minutes at room temperature. Platelets were then stimulated with increasing concentrations of reported P2Y_1_ receptor ligands and added to the top chamber of a 96-well transwell plate, with 0/30nM fMLP in the bottom chamber. After 90 minutes at 37°C, platelets in the bottom chamber were quantified and normalised to negative controls to give the chemotactic index (CI). (**A-B**) ADP-induced platelet chemotaxis following P2Y_1_ or P2Y_12_ receptor inhibition, respectively. (**C-D**) NAD. (**E-F**) ADP-ribose. (**G-H**) Up4A. Data: Mean ± SEM. n = 8 per group. One-way ANOVA with Tukey’s multiple comparisons. ^**^P<0.01, ^***^P<0.001, ^****^P<0.0001 versus in the presence of no nucleotide (column D). ^#^P<0.05, ^##^P<0.01, ^###^P<0.001, ^####^P<0.0001 versus positive control (column E).

We next assessed whether NAD^+^, ADP-ribose, and Up4A-P2Y_1_ -induced platelet chemotaxis was dependent on RhoA and Rac1 signalling events, as with ADP-induced platelet chemotaxis, rather than via activation of the canonical PLC signalling pathway required for Ca^2+^mobilization during P2Y_1_ induced platelet aggregation. Platelets were incubated in the presence of 0.1, 1, and 10 µM U73122 (PLC inhibitor), NSC23766 (Rac1 inhibitor), or GSK429286 (ROCK inhibitor) before stimulation with endogenous nucleotides, and chemotaxis was induced by fMLP. Incubation with NSC23766 had no effect on platelet chemotaxis induced by ADP, NAD^+^, ADP-ribose, or Up4A (**Figure 4A, 4D, 4G, 4J**). However, chemotaxis was significantly suppressed in the presence of increasing concentrations of NSC23766 (**Figure 4B, 4E, 4H, 4K**) and GSK429286 (**Figure 4C, 4F, 4I, 4L**). Thus, the endogenous nucleotides stimulate non-canonical P2Y_1_ signalling pathways (Rac1, and RhoA) to induce platelet chemotaxis.

**Figure 4.**
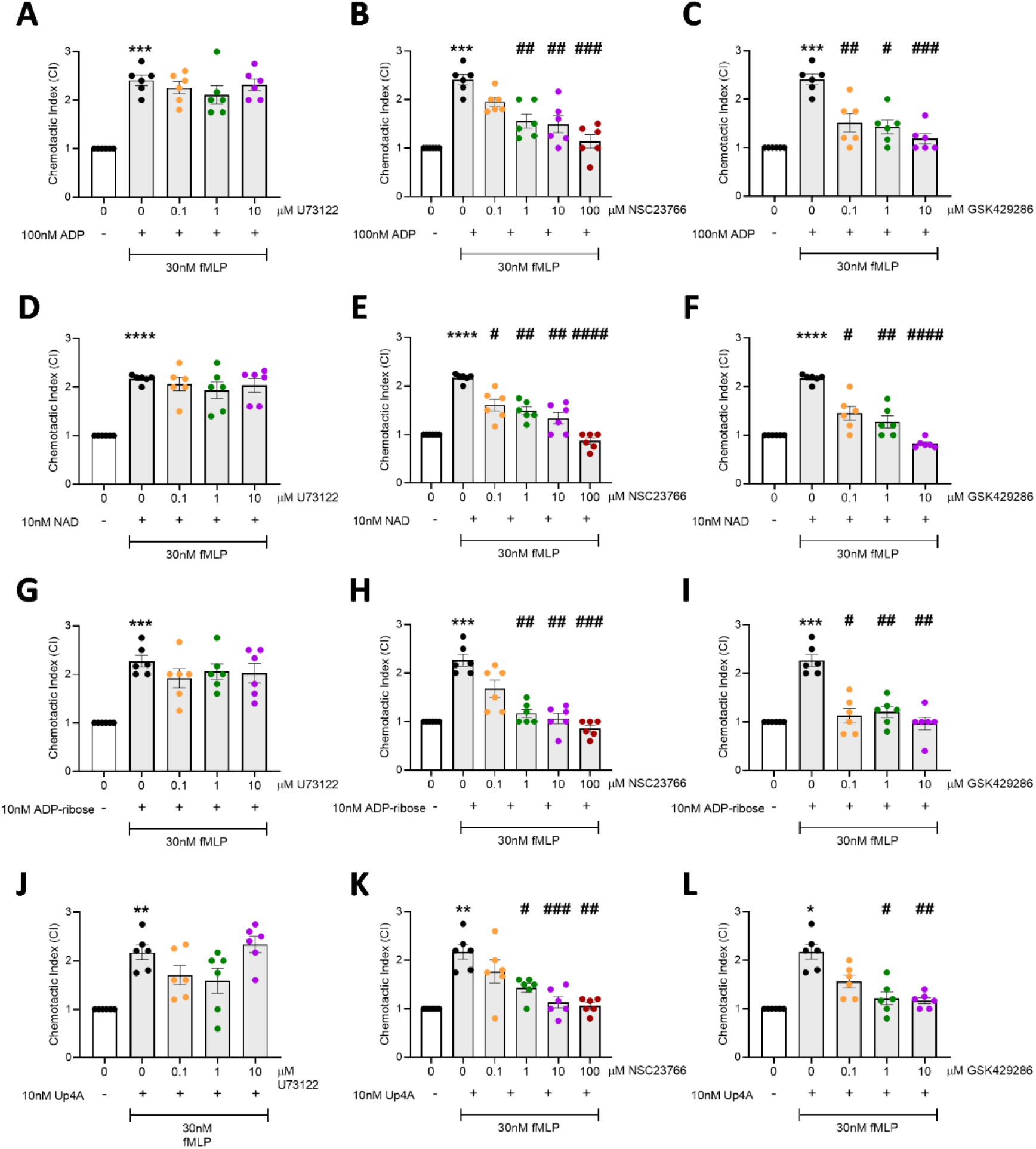
Rac1 and RhoA (ROCK) stimulation are necessary for P2Y_1_ ligand induced platelet chemotaxis. Washed platelets (5×10^7^/ml) were incubated with specific inhibitors for either PLC (U73122), Rac1 (NSC23766) or ROCK (GSK429286) for 10 minutes at room temperature. Platelets were then stimulated with increasing concentrations of endogenous P2Y_1_ receptor ligands and added to the top chamber of a 96-well transwell plate, with 0/30nM fMLP in the bottom chamber. After 90 minutes at 37°C, platelets in the bottom chamber were quantified and normalised to negative controls to give the chemotactic index (CI). (**A-C**) ADP-induced platelet chemotaxis following inhibition of PLC, Rac1 or ROCK, respectively. (**D-F**) NAD. (**G-I**) ADP-ribose. (**J-L**) Up4A. Data: Mean ± SEM. n = 6 per group. One-way ANOVA with Dunnett’s multiple comparisons. ^*^P<0.05, ^**^P<0.01, ^***^P<0.001, ^****^P<0.0001 versus negative control (column A). ^#^P<0.05, ^##^P<0.01, ^###^P<0.001, ^####^P<0.0001 versus positive control (column B).

### In silico molecular docking analyses of endogenous nucleotides to P2Y1 reveal different patterns of interaction compared to the non-biased agonist ADP

The functional studies described above revealed various endogenous nucleotides were able to activate platelets via P2Y_1_ receptors in a distinct manner compared to the cognate ligand ADP, which is able to activate platelets via canonical signalling pathways (PLC) to elicit aggregation, and alternative signalling pathways (RhoA, Rac1) to elicit motility (chemotaxis). NAD^+^, ADP-ribose, and Up4A, however, were only able to stimulate platelet chemotaxis, suggesting activation by alternative signalling pathways to aggregation. In order to better understand why these differences occurred between endogenous ligands, we performed a molecular docking analysis to compare their interaction with the P2Y_1_ receptor, with that of ADP. Using ChemPLP score, the binding energies of these endogenous ligands were investigated. The cognate P2Y_1_ receptor agonist, ADP, was found to have a ChemPLP score of 68.26, compared to 67.50 for Ap3A, 69.41 for NAD, 60.76 for ADP-ribose, and 66.26 for Up4A. Thus, alternative P2Y_1_ receptor ligands appeared to occupy the binding pocket of the P2Y_1_ receptor with a similar affinity to the cognate agonist, ADP, shown through comparable GOLD ChemPLP scores.

To understand the differences in the observed *in vitro* platelet function downstream of P2Y_1_ receptor activation by various ligands, the amino acids involved in these interactions were visualised *in silico* **(Figure 5A)** and compared **(Figure 5B)**. Of note, for the P2Y_1_ receptor, ADP was the only ligand found to interact with the hydrophilic amino acid, THR-206. In comparison, all other P2Y_1_ receptor ligands interacted with the positively charged amino acid, ARG-195, but ADP did not **(Figure 5B)**. Thus, unique patterns of amino acid interaction distinguish the non-biased P2Y1 agonist ADP to those with endogenous nucleotides (NAD^+^, ADP-ribose, and Up4A) that demonstrated biased agonist properties with respect to P2Y_1_-dependent platelet function, and this selective interaction with different amino acids within the binding pocket might play an important role in the observed bias for some endogenous ligands. The 3D interaction figures of the ligands with the P2Y1 receptor further reinforces this observation, as despite having similar ChemPLP scores, ADP sits deeper into the binding pocket of the P2Y1 receptor compared to other ligands which gives rise to differential amino acid interaction (**Figure 5C**).

**Figure 5.**
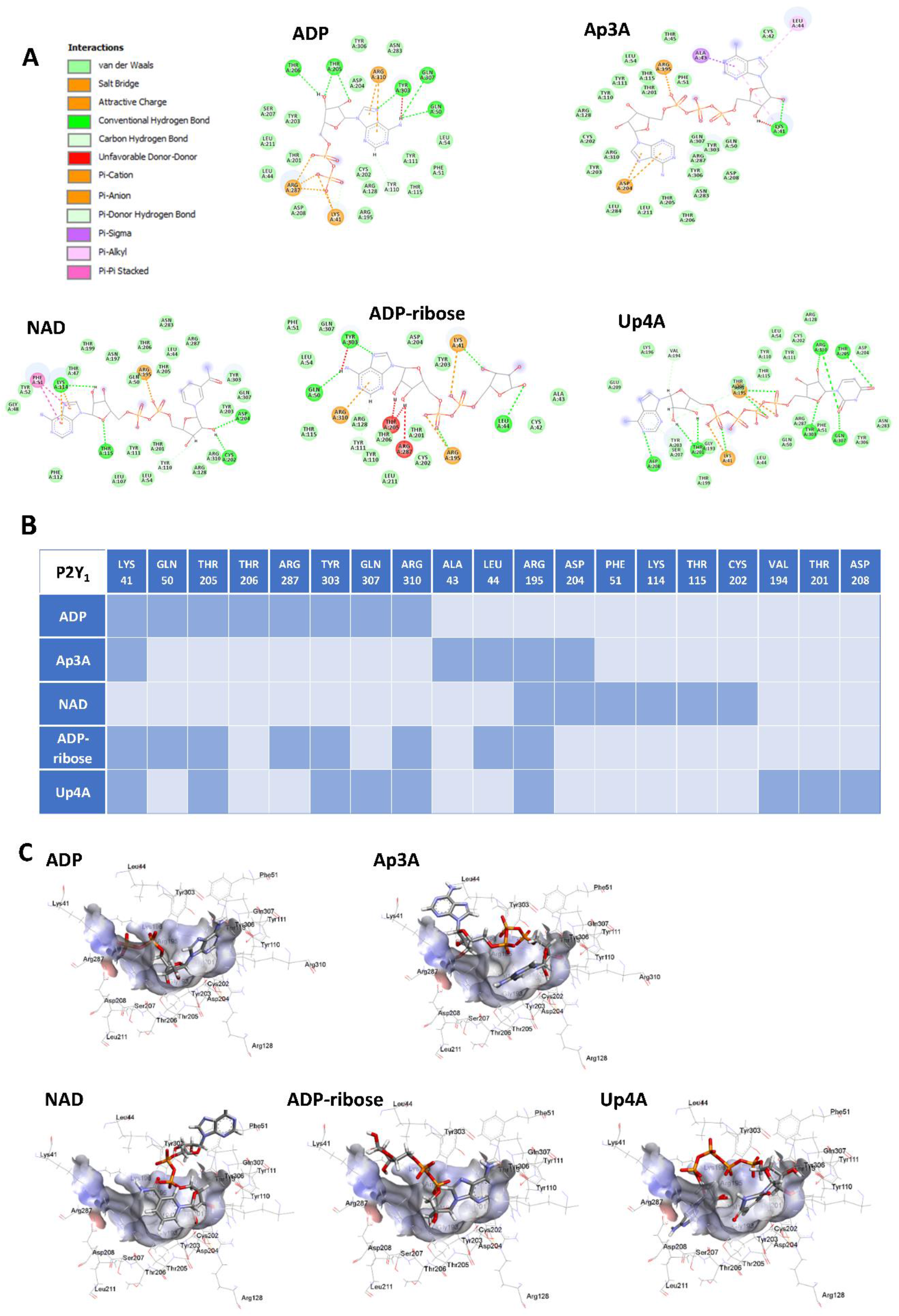
Computer modelling of endogenous P2Y_1_ receptor ligands reveal unique amino acid interactions for non-cognate ligands compared to ADP. 2D interaction diagrams showing how endogenous ligands bind to amino acids within the P2Y_1_ Receptor (**A**), and comparisons (**B**), identifying common versus unique contacts. 3D interaction diagram showing the relative position of the endogenous ligands within the P2Y1 binding pocket (**C**).

## Discussion

Although platelets were once thought of merely as cell fragments involved in aggregation, they are now widely appreciated as critical components of both haemostasis, host defence and inflammation. Evidence also suggests these different functions may be distinctly mediated. Our group have previously shown that inflammatory actions of platelets occurs downstream of the platelet P2Y_1_ receptor, in the absence of P2Y_12_ receptor activation (Amison *et al*., 2015; 2017; 2018). With this observation in mind, here we show differential effects of various endogenous nucleotides able to act as agonists at purinergic receptors on haemostatic and inflammatory functions of platelets *in vitro*.

In this study, we have used *in vitro* platelet functional assays to understand how platelet P2Y receptors mediate the dichotomy of platelet activation. We have utilised well-established *in vitro* assays of platelet aggregation, and fibrinogen binding, using both stimulated PRP or washed platelets to investigate the haemostatic actions of endogenous nucleotides. In addition, we have also investigated the ability of these endogenous nucleotides to induce platelet chemotaxis as an *in vitro* model of one of the inflammatory actions of platelets given the evidence that they can undergo extravascular migration into various tissues (Pitchford *et al*., 2008; Kraemer *et al*., 2010; Shah *et al*., 2021; Cleary *et al*., 2020). We have confirmed that platelets can undergo chemotaxis towards the robust chemotactic agent, fMLP as previously described (Czapiga *et al*., 2005; Amison *et al*., 2018) in the presence of ADP, the cognate purinergic P2Y_1_ and P2Y_12_ receptor agonist.

We have also shown that the endogenous nucleotide Ap3A also stimulates *in vitro* platelet aggregation. However, it has previously been shown that Ap3A itself is incapable of triggering platelet aggregation and that this effect is actually due to inherent hydrolase activity within plasma that converts Ap3A to ADP, which is then able to elicit an aggregatory response (Lüthje and Ogilvie, 1984; Lüthje, Baringer and Ogilvie, 1985). Since Ap3A is found in platelet dense granules and is released upon activation (Lüthje and Ogilvie, 1983), it is interesting to speculate its function. It may be that Ap3A acts as a competitive inhibitor to decrease ADP-induced platelet aggregation or, conversely, may act to sustain the aggregatory signal as it is hydrolysed to ADP. In contrast, none of the other endogenous nucleotides investigated (NAD^+^, ADP-ribose and Up4A) were found to exhibit any platelet aggregatory activity.

Interestingly, we have shown for the first time however, that along with ADP, the other endogenous P2Y agonists, NAD^+^, ADP-ribose and Up4A, are all able to elicit *in vitro* platelet chemotaxis towards fMLP via activation of P2Y_1_ receptors. When performing a concentration response to increasing ligand concentrations, a bell-shaped curved was observed. The reasons for this trend are not completely understood, but it could be that, at higher concentrations, the platelet P2Y1 receptor undergoes desensitisation and internalisation following ligand stimulation (Baurand *et al*., 2005; Hardy *et al*., 2005). Alternatively, high ligand concentrations might act as a self-regulatory mechanism to supress further platelet activation, since interaction between P2YR and adenosine (released or metabolised) signalling pathways (that inhibit both haemostasis and inflammatory events) has been reported (Shih et al., 2021; Layland et al., 2014). Like ADP, NAD^+^, ADP-ribose and Up4A induced chemotaxis via Rac1 and RhoA-dependent signalling pathways. Thus, their inability to promote platelet aggregation or fibrinogen binding, demonstrated an absence of activation of the P2Y_1_ canonical PLC signalling pathway, suggesting that these endogenous nucleotides exhibit biased agonist properties, with functional selectivity of platelet activation involved in inflammation (Kenakin 2012).

*In silico* analysis to compare the ability of NAD^+^, ADP-ribose and Up4A to interact with the P2Y_1_ receptor demonstrated that these endogenous ligands interacted with the receptor with similar affinities to ADP, but that their relative position within the binding pocket were slightly different. Furthermore, these experiments showed that the endogenous nucleotides showed interactions with amino acids that differed to those recognised by ADP, providing a potential mechanism by which these biased properties occur.

The relevance of these biased differences in platelet activation induced by different endogenous nucleotides acting on P2Y receptors leading to chemotaxis (ie a non-thrombotic action) without inducing aggregation or fibrinogen binding have not yet been investigated *in vivo*. However, it is likely that platelet function is influenced by an integrated response to the mixed extracellular milieu of nucleotides (a ‘nucleotide halo’) that can act with distinct properties via the same receptor type, or other purinergic receptors expressed on platelets and activated by distinct nucleotides (for example ATP, UDP-glucose), released during trauma, to dictate an inflammatory set of functions, as opposed to haemostatic responses. Greater understanding of this biased agonism pathway may lead to the development of novel pharmacological strategies to target specific platelet functions applicable to inflammation and host defence (Pitchford *et al*., 2019).

## Funding Acknowledgement

This research was funded by a Medical Research Council Doctoral Training Grant (MRC-DTP), awarded to Dr Simon Pitchford to fund Kate Arkless (MR/N013700/1); and MRC project grant (MR/T015845/1) awarded to Dr Simon Pitchford.

