## Supplementary figures and images for "Stimulation of platelet P2Y_1_ receptors by different endogenous nucleotides leads to functional selectivity via biased signalling"

### Supplementary Data

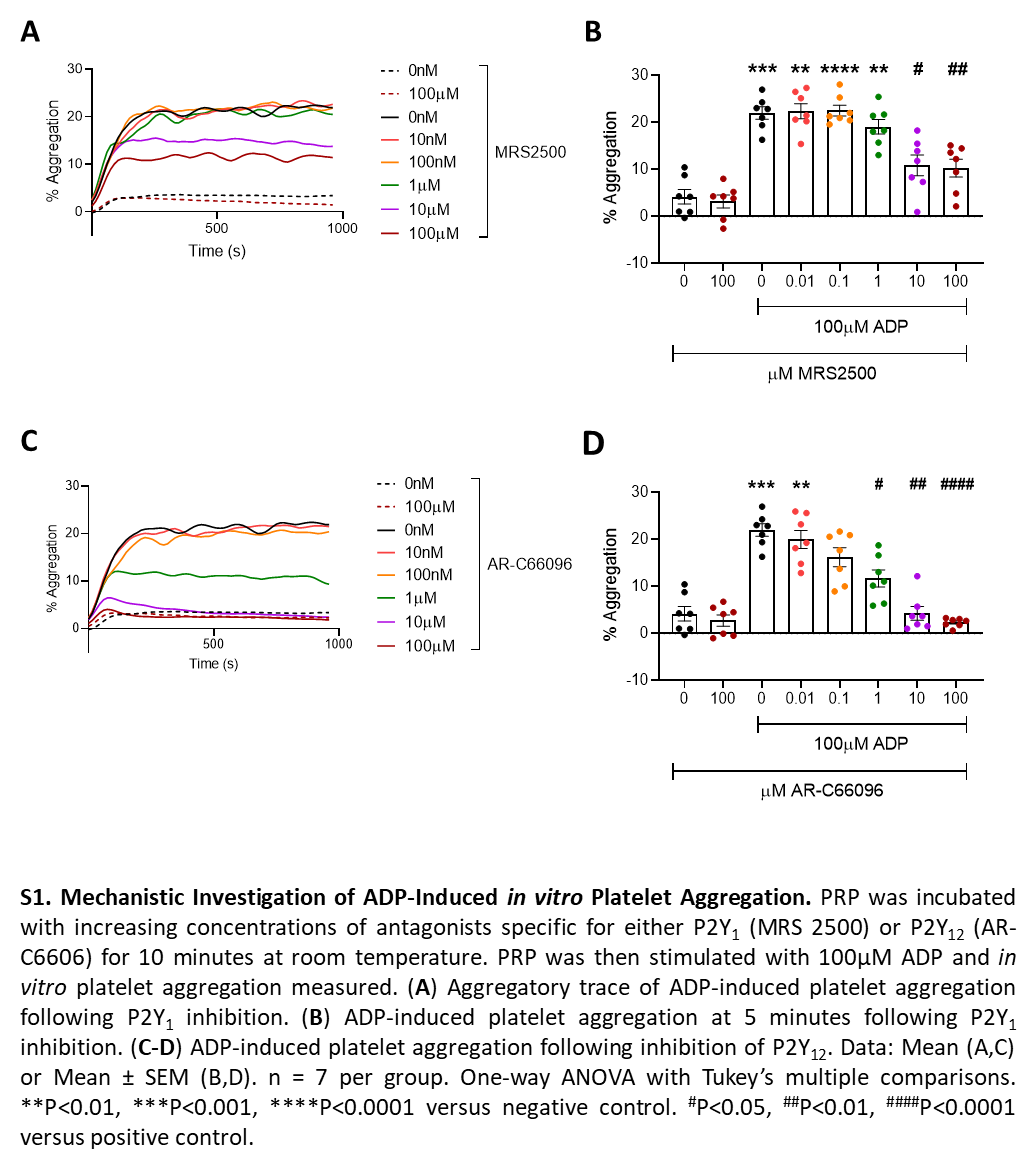

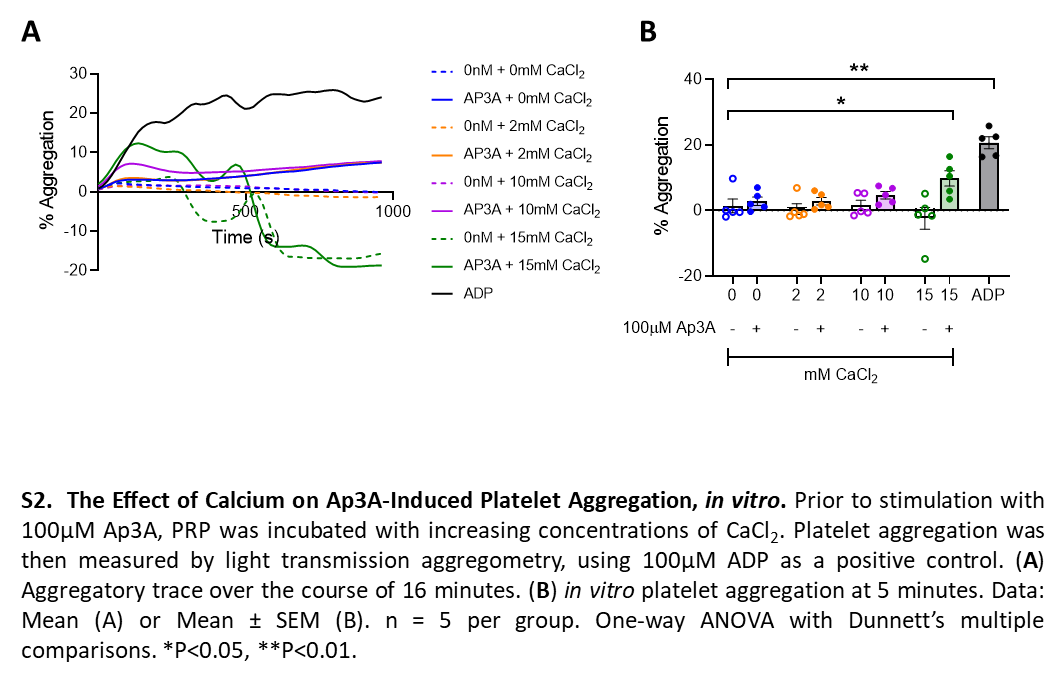
